# Spatiotemporal dynamics and ossification of podoplanin-positive cells in the developing femur

**DOI:** 10.1101/2024.05.15.593859

**Authors:** Hinako Notoh, Nagaharu Tsukiji, Nobuaki Suzuki, Atsuo Suzuki, Shuichi Okamoto, Takeshi Kanematsu, Akira Katsumi, Rikuto Nara, Ayuka Kamata, Tetsuhito Kojima, Tadashi Matsushita, Katsue Suzuki-Inoue, Shogo Tamura

## Abstract

In vertebral long bones, such as the femur, bone formation involves endochondral ossification. Endochondral ossification first occurs in the central region of the fetal diaphysis, the primary ossification center. Podoplanin (PDPN) is a transmembrane mucin-like glycoprotein, and PDPN-positive cells play key roles in various organ/tissue development. In adult mice, osteolineage PDPN-positive cells are associated with femoral microarchitecture. However, specific roles of PDPN in fetal bone development remain unclear. Therefore, in this study, we aimed to investigate the spatiotemporal dynamics and physiological functions of PDPN-positive cells during fetal femur development. In the fetal femur, PDPN-positive cells first emerged in the primitive cortical bone, termed the bone collar, concurrently with primary ossification center initiation. Several PDPN-positive cells in the bone collar migrated to the marrow cavity and populated the metaphyseal trabecular bone. Most PDPN-positive cells in both the bone collar and trabeculae exhibited osteolineage features, such as osterix expression. *Pdpn* knockout fetuses exhibited abnormal recruitment of osterix-positive cells and mineral deposition in the dorsal bone collar. Overall, our results suggest that PDPN-positive cells constitute a spatially regulated osteolineage population that contributes to coordinated fetal femur development.

## Introduction

In vertebrates, bone formation occurs via two distinct developmental processes during the fetal and neonatal stages: Intramembranous and endochondral ossification ^1,2^. Intramembranous ossification is the developmental process of flat bones, such as the skull and clavicle. In this process, bone is directly differentiated from the mesenchymal stem cells (MSCs) or mesenchymal stem/progenitor cells in the soft connective tissue, without any intermediate cartilage. In contrast, endochondral ossification is responsible for the development of most bones in the body, except flat bones. Endochondral ossification is a highly regulated stepwise process involving the coordinated actions of multiple cell types.

Endochondral ossification begins with the formation of the avascular primitive cartilage originating from condensed MSCs ^3–8^. Chondrocytes in the center of primitive cartilage further differentiate and mature into hypertrophic chondrocytes, which express several angiogenic factors, such as the fibroblast growth factor-2, matrix metalloproteinase-9, and vascular endothelial growth factor. These angiogenic factors induce the invasion of the surrounding perichondral vasculature, mesenchymal stromal cells, and skeletal cells, including bone-forming osteoblasts and bone-resorbing osteoclasts/chondroclasts, into the cartilage, facilitating the replacement of cartilage with bone and initiating bone marrow development^3,5,7^. In long bones, such as the femur, endochondral ossification occurs first in the central region of the fetal diaphysis, the primary ossification center (POC), and later in the neonatal epiphysis, the secondary ossification center ^6,8^. In mice, around embryonic day (E)-14.5, POC formation is accompanied by the development of a “bone collar,” a structure formed by osteoblasts differentiated from the perichondrium. This fetal bone collar forms the “cortical bone” after birth ^9^.

Podoplanin (PDPN) ^10^, also known as E11 ^11^, gp38 ^12^, or T1a ^13^, is a transmembrane mucin-like glycoprotein expressed in multiple stromal cell types in both the developmental and adult tissues ^14^. PDPN-positive cells are found in various organs, such as the brain, lungs, lymphatic vessels, and heart, where they play key roles in organogenesis ^14^. PDPN is potentially associated with long bone formation. In adult mice, PDPN is expressed in the periosteal osteoblasts and early differentiated osteocytes within the diaphyseal cortical and metaphyseal trabecular bones of the femur ^15,16^. *Pdpn* conditional knockout (KO) in the osteogenic lineage or KO of the fibroblast growth factor-2, a *Pdpn* upstream regulator, causes mild abnormalities in the cortical bone microarchitecture, without affecting the trabecular bone ^17–20^. However, specific roles of PDPN in fetal bone development remain unclear. To better understand PDPN-associated bone development, this study aimed to assess the physiological functions of PDPN-positive cells during fetal femur development. Here, we report the spatiotemporal dynamics of PDPN-positive osteolineage cells within the developing femur and their functional roles in bone collar formation.

## Results

### Spatiotemporal emergence of PDPN-positive cells during fetal femur development

First, we conducted histological analysis to examine the spatiotemporal distribution of PDPN-positive cells within the POC-associated diaphyseal and metaphyseal regions of the femur. According to the nomenclature used by Maes et al. ^9^, diaphyseal bone is referred to as the “bone collar” during embryonic/fetal stages and “cortical bone” post-birth (Fig. 1A). Trabecular bone is primarily found in the epiphyses and metaphyses but almost absent in the mid-diaphysis. As POC formation and diaphyseal marrow development begin on E14.5 ^5,9^, we examined the fetal femur daily from E13.5–E18.5 and traced PDPN-positive cells within the developing bone and marrow (Fig. 1B). On E13.5 and E14.5, the femur consisted of cartilage tissue formed by chondrocytes and surrounded by a thin outer layer, called the perichondrium (Fig. 1C and D). Chondrocytes at the center of the diaphysis had increased in size and differentiated into hypertrophic chondrocytes by E14.5. No blood vessels or bone-forming cells, including PDPN-positive cells, were observed in the cartilage at this stage (Fig. 1C and D). On E15.5, CD31-positive blood vessels began to invade the cartilage template and some PDPN-positive cells were observed at the bone collar (Fig. 1E). A bone marrow cavity was formed and PDPN-positive cells expanded within the bone collar on E16.5 (Fig. 1F). PDPN-positive cells were distinctly observed in the bone collar and primitive bone marrow on E17.5 (Fig. 1G), with the number of PDPN-positive cells in the bone collar being higher than that observed on E16.5. Notably, several PDPN-positive cells appeared to migrate from the bone collar to the bone marrow (Fig. 1G, arrowheads). On E18.5, PDPN-positive cells were abundantly distributed within the bone collar and along the primitive trabecular bone (Fig. 1H). In the trabecular bone, PDPN-positive cells were restricted to the marrow cavity-side (Fig. 1H-i and ii; marrow cavity-side of the trabecular bone is indicated by the yellow dotted line). These observations suggest that PDPN-positive cells emerge and expand within the bone collar during POC formation, invade the primitive cartilage, and migrate to the marrow cavity-side of the trabecular bone by E17.5. Moreover, PDPN-positive cells play key roles in bone collar and/or trabecular bone formation due to colocalization.

**Figure 1.**
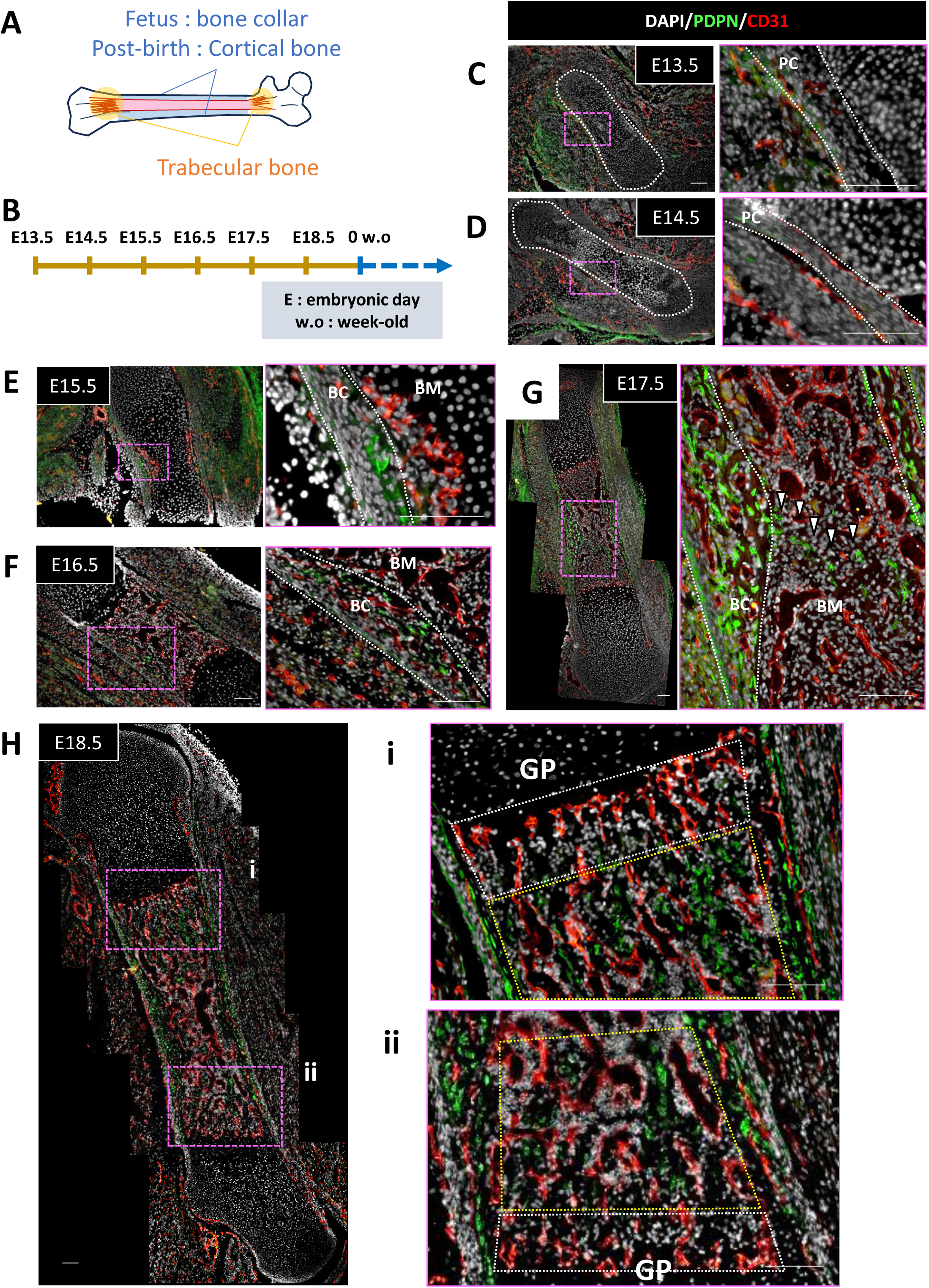
Spatiotemporal dynamics of podoplanin (PDPN)-positive cells during fetal femur development. (A) Schematic illustration indicating the anatomical regions of the femur. For clarity, diaphyseal bone is referred to as the “bone collar” during the embryonic/fetal stages and “cortical bone” post-birth. (B) Schematic diagram of the experimental design, including the analyzed developmental time points. (C–H) Representative immunohistochemical images of the femur on embryonic day (E)-13.5 (C), E14.5 (D), E15.5 (E), E16.5 (F), E17.5 (G), and E18.5 (H). PDPN-positive cells are shown in green. CD31-positive endothelial cell signals, indicating blood vessels, are indicated in red. Grid-lined areas indicate the primitive cartilage. Arrowheads indicate the PDPN-positive cells migrating to the bone marrow cavity. BC, bone collar; BM, bone marrow; GP, growth plate. Scale bar, 100 μm.

### PDPN-positive cell distribution aligns with the onset of cortical and trabecular ossification

To determine the roles of PDPN-positive cells in bone formation in the developing femur, we examined the histological progression of fetal femur from E16.5– E18.5 via hematoxylin-eosin (HE), alcian blue, and von Kossa staining (Fig. 2). Alcian blue visualizes the cartilage by staining with acidic mucopolysaccharides, and von Kossa detects the mineral deposits indicative of bone. To assess the spatial relationship between PDPN-positive cells and osteogenic components, we conducted immunohistochemical (IHC) analysis of PDPN-positive cells in combination with osterix (OSX)-positive cells representing the immature osteoblasts (Fig. 3).

**Figure 2.**
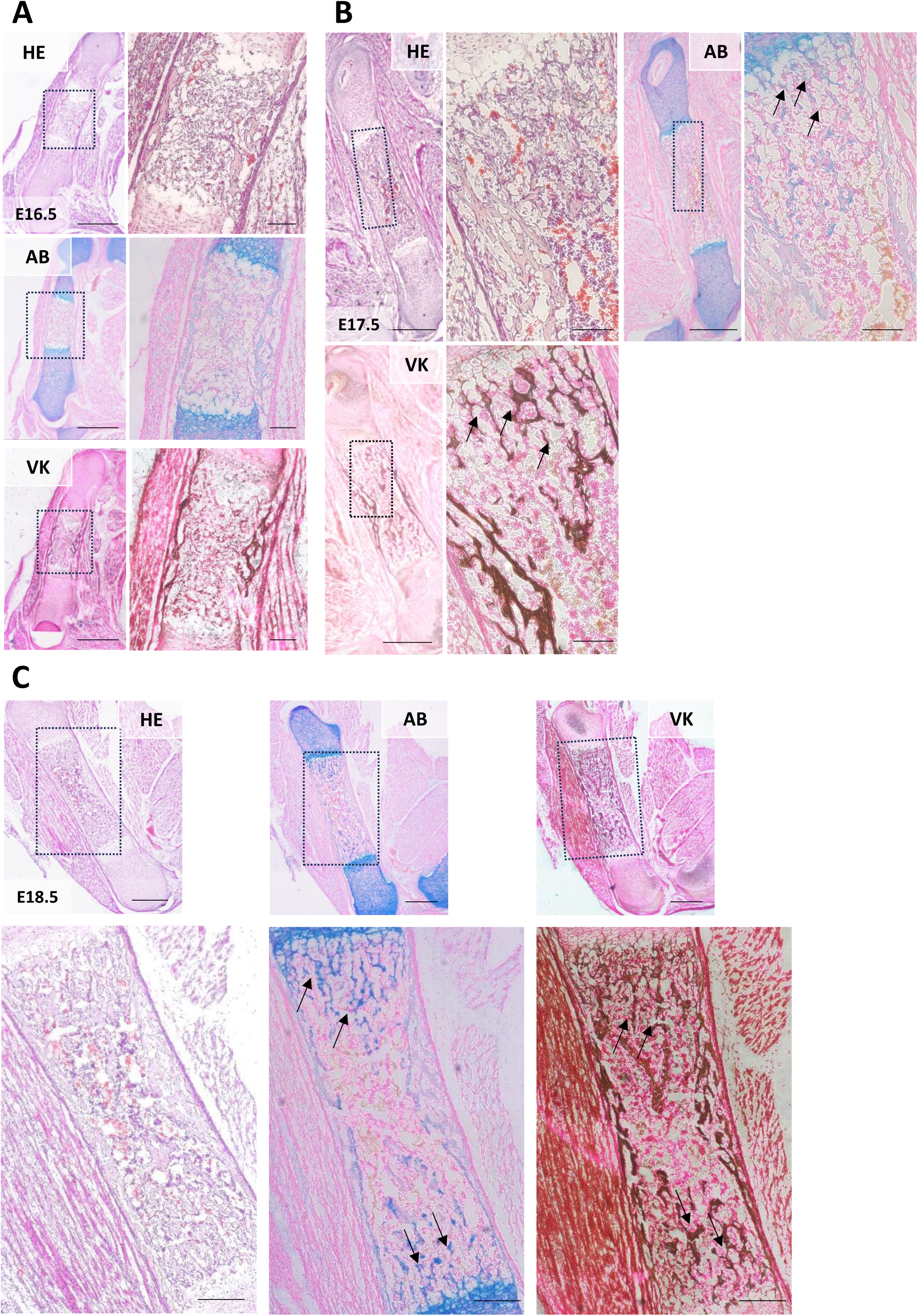
Histological progression of fetal femur development. (A–C) Representative images of hematoxylin–eosin (HE), alcian blue (AB), and von Kossa (VK) staining of the femur on E16.5 (A), E17.5 (B), and E18.5 (C). Alcian blue staining visualized the cartilage by staining with acidic mucopolysaccharides. Von Kossa staining detected the mineral deposits indicative of bone. Black arrows indicate the trabecular bone structure. Scale bar, 500 μm.

**Figure 3.**
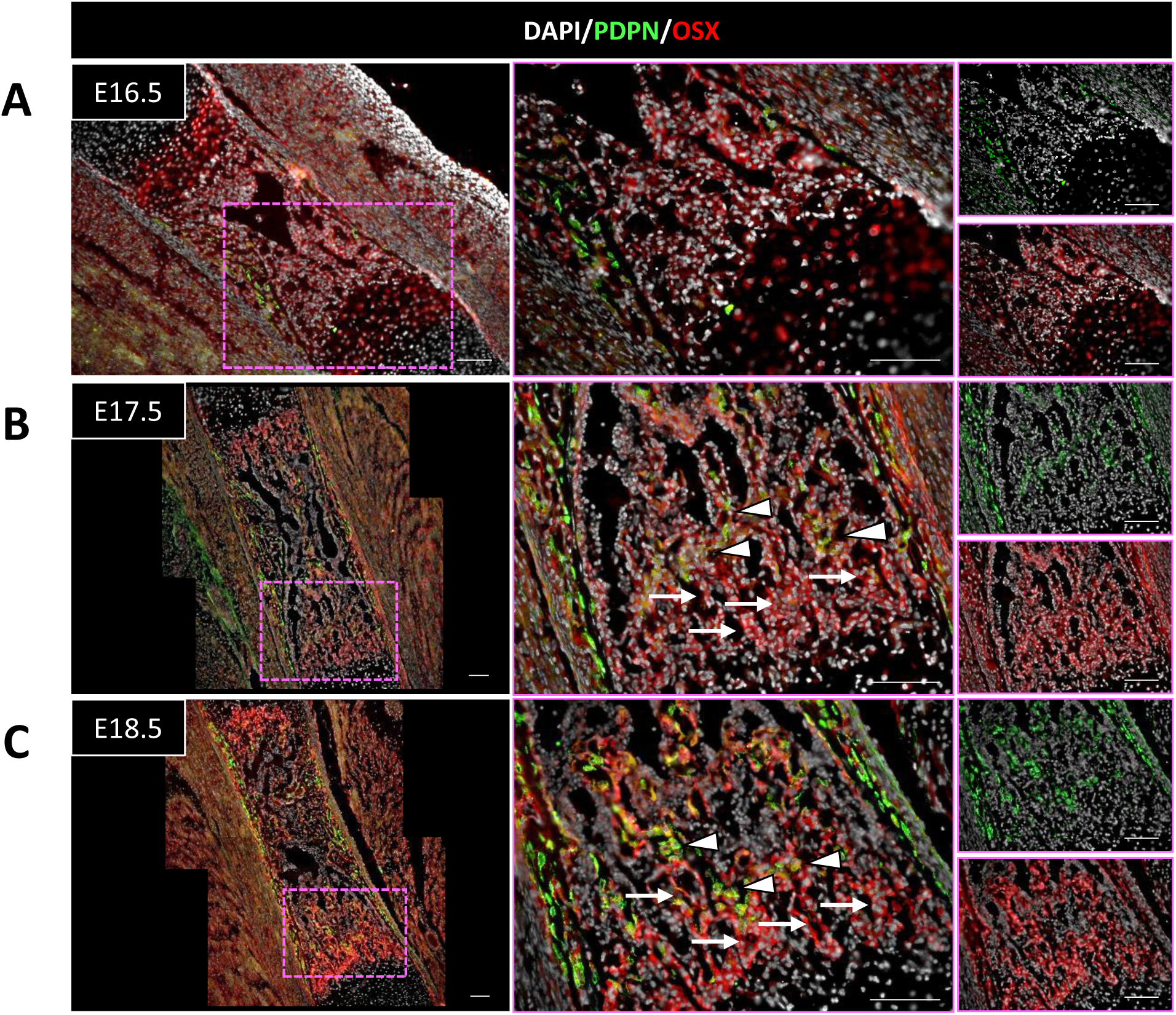
Spatiotemporal association of PDPN-positive cells with the osteogenic components of the developing fetal femur. Representative immunohistochemical images targeting PDPN/osterix (OSX) of the femur on E16.5 (A), E17.5 (B), and E18.5 (C). PDPN-positive cells are shown in green. OSX-positive cells, indicating osteogenic cells, are shown in red. White arrows indicate the trabecular bone structures, and white arrowheads indicate the PDPN-positive cells. Scale bar, 100 μm.

On E16.5, the bone collar was stained with von Kossa, and trabecular region was weakly stained with alcian blue and von Kossa (Fig. 2A). On E17.5, similar to the bone collar, trabecular region was clearly stained with both alcian blue and von Kossa (Fig. 2B). Von Kossa-positive branched and thin structures indicated that the trabecular bone was more developed on E18.5 than on E17.5 (Fig. 2C, arrows). Subsequently, we performed IHC staining of PDPN-positive cells and bone-associated markers to determine whether PDPN-positive cells correspond to bone collar and trabecular bone formation.

In IHC analysis on E16.5, OSX-positive cells were detected both in the bone collar and primitive marrow (Fig. 3A). At this point, PDPN-positive cells were restrictedly observed in the bone collar. Trabecular bone structures, including OSX-positive cells, were evident (Fig. 3B, arrows), and PDPN-positive cells were observed both in the bone collar and marrow cavity-side of the trabecular bone on E17.5 (Fig. 3B, arrowheads). Notably, PDPN-positive cells were more prominent in the trabecular bone on E18.5 than on E17.5 (Fig. 3C). These results suggest that the cellular dynamics of PDPN-positive cells are associated with femoral bone development. Particularly, PDPN-positive cell migration to the marrow-facing trabecular region coincides with the onset of trabecular bone formation between E16.5 and E17.5 and is enhanced by E18.5.

### Neonatal redistribution of PDPN-positive cells in the femur

During fetal bone and marrow development, PDPN-positive cells were localized in the bone collar and marrow cavity-side of the trabecular bone. To investigate the ways in which the tissue distribution of PDPN-positive cells changes after birth, we analyzed the postnatal femurs of 0–3 weeks old (w.o.) mice (Fig. 4). At 0 and 1 w.o., PDPN-positive cells were abundantly distributed throughout the cortical and trabecular bones (Fig. 4A). However, numbers of PDPN-positive cells in the cortical and trabecular bones were decreased and their distribution became more restricted, predominantly localized to the periosteum, endosteum, and trabecular bone regions, in 2 and 3 w.o. mice (Fig. 4B and D). Quantification of PDPN-positive cells revealed cortical and trabecular bone transition during postnatal development (Fig. 4C and E). In the cortical bone, including the periosteum and endosteum, number of PDPN-positive cells decreased from 0 w.o. to 1 w.o., with a significant reduction observed between 1w.o. and 2 w.o. (number of PDPN-positive cells in the cortical bone: 2177.15 ± 540.55, 1328.89 ± 319.55, and 980.12 ± 267.30 cells at weeks 0, 1, and 2 w.o., respectively; Fig. 4B and C). From 2 w.o. onward, PDPN-positive cells in the cortical bone were mainly localized to the periosteum and limited regions of the endosteum. In the trabecular bone, number of PDPN-positive cells remained constant between 0 w.o. and 1 w.o., but significantly decreased at 2 w.o. (number of PDPN-positive cells in the metaphysis: 666.22 ± 153.37, 590.30 ± 75.38, and 363.31 ± 60.71 cells at 0, 1, and 2 w.o., respectively; Fig. 4D and E). These results suggest that PDPN-positive cells are widely distributed in the neonatal femur, with their localization becoming progressively restricted to specific regions (periosteum, endosteum, and trabecular bone) during femur development.

**Figure 4.**
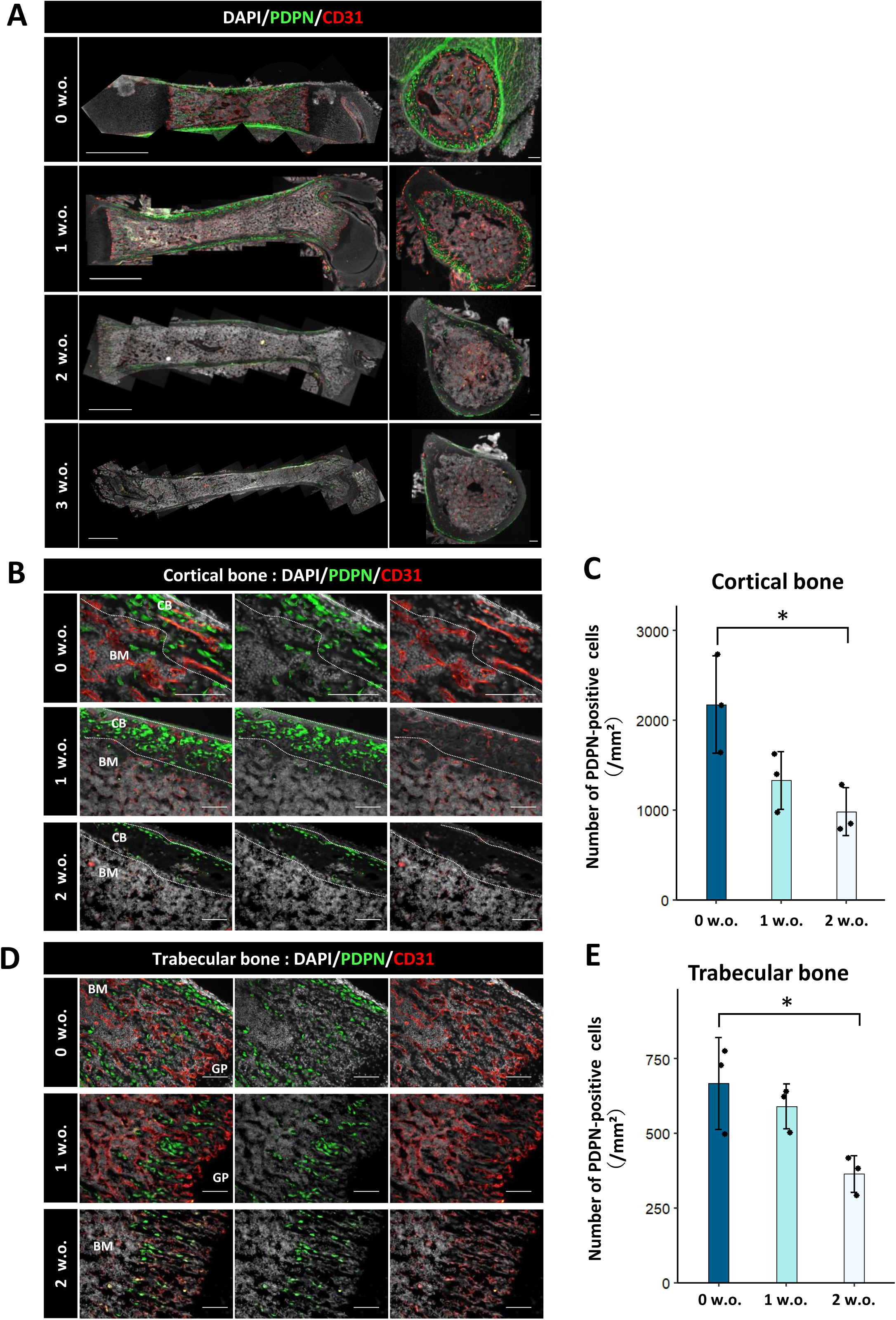
Localized distribution of PDPN-positive cells in the postnatal femur. (A) Representative immunohistochemical images of the femur in 0–3 weeks old (w.o.) mice. (B) Representative images of the cortical bone region from 0 to 2 w.o. (C) Quantification of PDPN-positive cells in the cortical bone region from 0 to 2 w.o. (D) Representative images of the trabecular bone region from 0 to 2 w.o. (E) Quantification of PDPN-positive cells in the trabecular bone region from 0 to 2 w.o. In panels A, B and D, PDPN-positive cells are shown in green. CD31-positive endothelial signals, indicating blood vessels, are shown in red. Scale bar, 100 μm. In panels C and E, data are represented as the mean ± standard deviation (SD; n = 3/group). Statistical comparisons were performed via one-way analysis of variance (ANOVA), followed by the Tukey–Kramer post-hoc test. \**p* < 0.05.

### PDPN-positive cells exhibit osteolineage characteristics in the bone collar/cortical bone and periosteum

As PDPN-positive cells were associated with femoral bone development, we further examined whether they exhibited osteolineage features during the fetal and neonatal stages. PDPN marks various osteogenic subsets, such as late-stage osteoblasts and early osteocytes in adult mice ^11,21–23^. However, specific roles of PDPN-positive cells in fetal and neonatal bone modeling remain unclear. Since immature osteoblasts, marked by OSX, are abundant in the fetal perichondrium and bone collar ^5,7,8^, we hypothesized that PDPN-positive cells in the fetal femur represent an osteolineage cell subpopulation. To verify this, we characterized PDPN-positive cells in the fetal/postnatal femur via IHC analysis using OSX as a marker of immature osteoblasts and osteocalcin (OCN) as a mature osteoblast marker (Fig. 5A).

**Figure 5.**
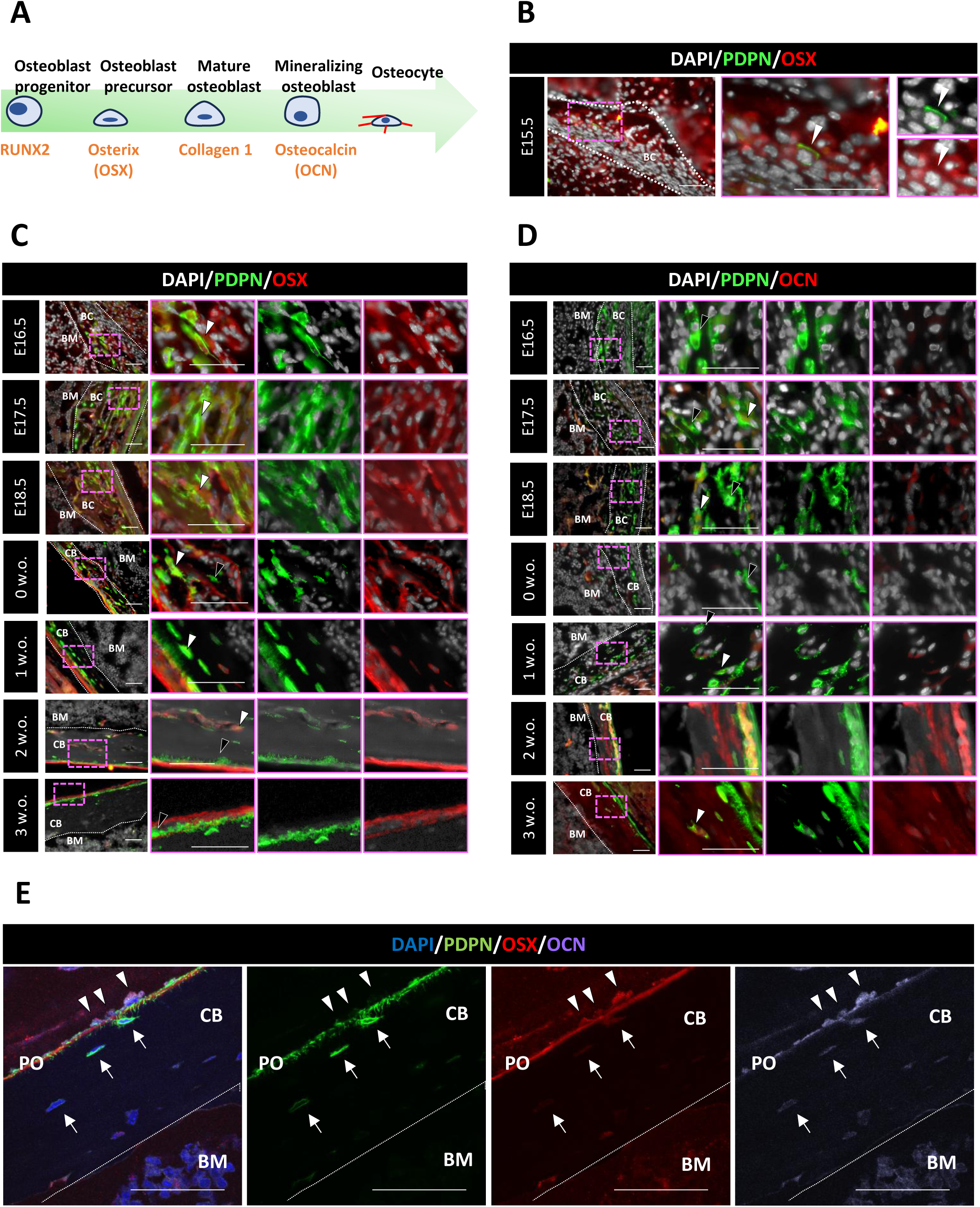
PDPN-positive cells in the developing bone collar/cortical bone and periosteum. (A) Schematic illustration of osteolineage differentiation, showing the representative cell markers at each stage. (B) Representative immunohistochemical images of the bone collar on E15.5. (C) Representative images of the bone collar from E16.5 to week 3 post-birth. In panels B and C, PDPN-positive cells are shown in green. OSX-positive cells (osteoblast precursors) are shown in red. White arrowheads indicate the OSX- and PDPN-positive cells. Black arrowheads indicate the OSX-negative PDPN-positive cells. Scale bar, 50 μm. (D) Representative images of the bone collar from E16.5 to 3w.o. PDPN-positive cells are shown in green. Osteocalcin (OCN)-positive cells (mineralizing osteoblasts and osteocytes) are shown in red. White arrowheads indicate the OCN- and PDPN-positive cells, and black arrowheads indicate the OCN-negative PDPN-positive cells. Scale bar, 50 μm. (E) Representative confocal microscopy images of the cortical bone at 3 w.o. PDPN-positive cells are shown in green. OSX and OCN are shown in red and cyan, respectively. White arrowheads indicate the layered PDPN-positive cells in the periosteum. White arrows indicate the sporadic PDPN-positive cells in the cortical bone. Scale bar, 50 μm.

First, we analyzed the fetal bone collar and postnatal cortical bone (Fig. 5B–E). In the IHC analysis for OSX, OSX-positive cells were observed on E15.5, and a small PDPN/OSX-positive cell population was found in the bone collar (Fig. 5B). Cells in the bone collar widely expressed OSX, and number of PDPN/OSX-positive cells increased from E16.5 to E18.5 (Fig. 5C). PDPN-positive, OSX-positive, and OSX-negative cells were detected in the cortical bone at 0–1 w.o. (Fig. 5C). By 2–3 w.o., a small number of PDPN-positive cells were detected in the cortical bone (Fig. 5C, filled arrowhead). In the periosteum, PDPN-positive cells were observed in a layered pattern with villous projections; however, whether the PDPN signals overlapped with the OSX signals remains unclear (Fig. 5C, white arrowhead). IHC analysis on E16.5 revealed that the PDPN-positive cells in the bone collar did not express OCN (E16.5; Fig. 5D). PDPN-positive, OCN-positive, and OCN-negative cells were observed from E17.5 to week 1 post-birth (Fig. 5D). From 2 w.o. onward, OCN was broadly detected throughout the cortical bone, and PDPN/OCN-positive cells were sporadically observed within the cortical bone (2 –3 w.o.; Fig. 5D). However, whether layered periosteal PDPN-positive cells with villous projections express OCN remains unclear.

To further resolve the osteolineage characteristics of PDPN-positive cells in the cortical bone, we performed high-resolution confocal imaging with triple immunostaining for PDPN, OSX, and OCN in the femur at 3 w.o. (Fig. 5E). Sporadic PDPN-positive cells in the cortical bone co-expressed OSX and OCN, representing late-stage osteoblasts that transitioned to early osteocytes (Fig. 5E, white arrows). In contrast, layered PDPN-positive cells in the periosteum exhibited compartmentalized marker expression, and their cell bodies were weakly positive for PDPN and positive for both OSX and OCN, whereas their extended pseudopods were strongly PDPN-positive but negative for OSX and OCN (Fig. 5E, white arrowheads). According to Nagai et al. ^16^, cells with villous projections and such expression patterns correspond to late-stage osteoblasts. Collectively, these results suggest that PDPN-positive cells initially develop into OSX-positive cells in the fetal bone collar and subsequently contribute to cortical bone development and modeling during postnatal growth.

### Heterogeneous osteolineage potential of PDPN-positive cells in the trabecular bone

Next, we investigated the characteristics of PDPN-positive cells in the trabecular bone from E16.5 to week 3 post-birth (Fig. 6). Consistent with the data shown in Fig. 3A–C, IHC analysis for PDPN and OSX revealed that the OSX-positive cells were broadly distributed in the primitive trabecular region from E16.5, followed by the emergence of PDPN-positive cells in the same region (E16.5–E18.5; Fig. 6A). From E17.5 to 0 w.o., PDPN- and OSX-positive cells were observed along the trabecular bone surface. From 1 w.o. onward, two PDPN-positive cell population types, OSX-positive and -negative populations, were detected on the trabecular surface (2–3 w.o.; Fig. 6A). Notably, IHC analysis for PDPN and OCN revealed no OCN-positive cells in the primitive trabecular region on E16.5 (Fig. 6B). From E17.5 onward, OCN expression became gradually evident in the trabecular bone-forming cells. A subset of PDPN-positive cells on the trabecular surface also showed OCN positivity during this period (E17.5 to 3 w.o.; Fig. 6B).

**Figure 6.**
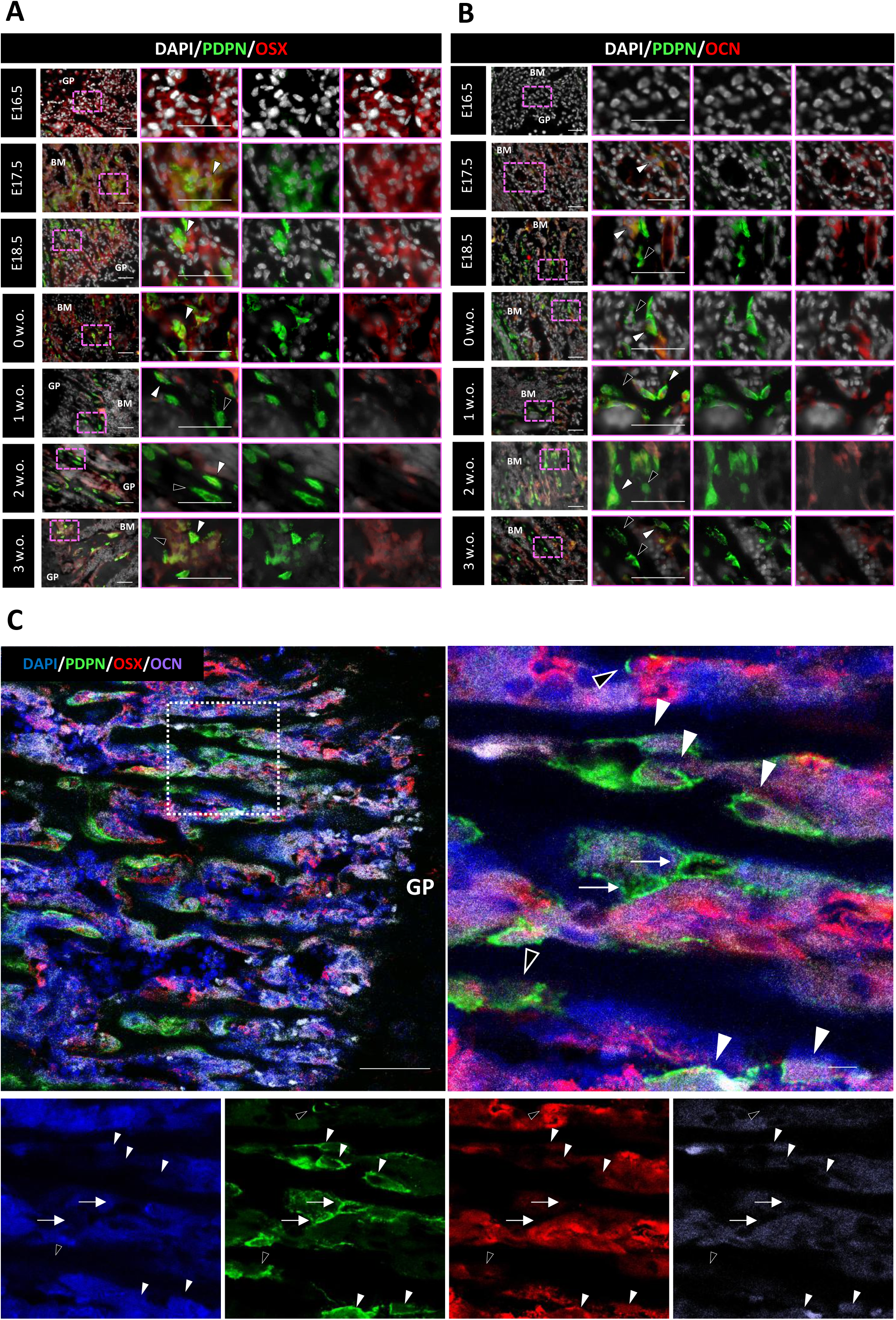
PDPN-positive cells in the developing trabecular region. (A) Representative immunohistochemical images of the trabecular bone from E16.5 to week 3 post-birth. PDPN-positive cells are shown in green. OSX-positive cells (osteoblast precursors) are shown in red. White arrowheads indicate the OSX- and PDPN-positive cells. Black arrowheads indicate the OSX-negative PDPN-positive cells. Scale bar, 50 μm. (B) Representative images of the trabecular region from E16.5 to 3w.o. PDPN-positive cells are shown in green. OCN-positive cells (mineralizing osteoblasts and osteocytes) are shown in red. White arrowheads indicate the OCN- and PDPN-positive cells. Black arrowheads indicate the OCN-negative PDPN-positive cells. Scale bar, 50 μm. (C) Representative confocal microscopy images of the trabecular bone at 3 w.o. PDPN-positive cells are shown in green. OSX and OCN are shown in red and cyan, respectively. White arrowheads indicate the PDPN-positive cells co-expressing OSX and OCN. Black arrowheads indicate the PDPN-positive cells co-expressing OSX, but not OCN. White arrows indicate the PDPN-positive cells lacking OSX and OCN expression. Scale bar, 100 μm.

To further examine the heterogeneity of PDPN-positive cells in this region, we performed high-resolution confocal imaging with triple immunostaining for PDPN, OSX, and OCN (Fig. 6C). Most PDPN-positive cells associated with the trabecular bone co-expressed OSX and OCN (Fig. 6C, white arrowheads), indicating osteolineage commitment. A small cell subset was OSX-positive but OCN-negative (Fig. 6C, black arrowheads), suggesting an immature osteoblast-like phenotype. In contrast, a minor population of PDPN-positive cells was negative for both markers (Fig. 6C, arrows), indicating the presence of non-osteolineage cells. Notably, no PDPN-positive/OSX-negative/OCN-positive cells were observed. These findings suggest that PDPN-positive cells infiltrating the primitive marrow during trabecular bone formation predominantly consist of osteolineage cells, but also include a minor cell population lacking the classical osteogenic markers.

### PDPN-positive cells possess a role for the proper mineralization of the fetal bone collar but dispensable for trabecular bone development

Considering the spatiotemporal and phenotypic associations between PDPN-positive cells and developing bone structures, we investigated whether PDPN is functionally necessary for proper femoral bone formation by comparing bone formation in *Pdpn* wild-type (WT) and KO mice (Figs. 7 and 8).

**Figure 7.**
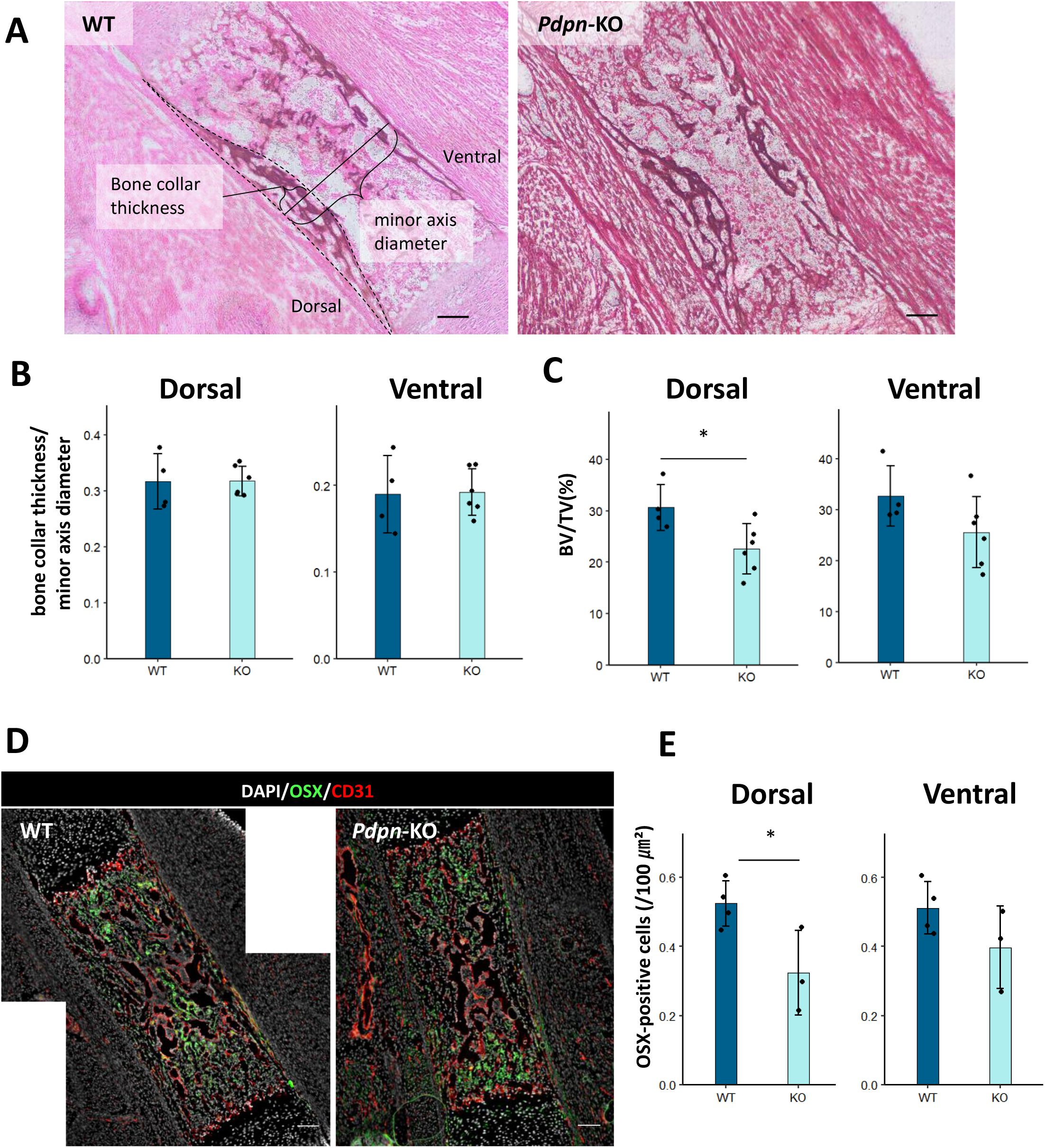
Bone collar osteogenesis in fetal *Pdpn* knockout (KO) mice. (A) Representative longitudinal images of the fetal femur in the *Pdpn* wild type (WT) and KO mice on E17.5. The sections were stained with von Kossa. Morphological parameters, such as bone collar thickness and minor axis diameter, are indicated. (B) Quantification of bone collar thickness in the *Pdpn* WT and KO E17.5 fetal femurs. To compensate for interindividual variability, bone collar thickness was normalized to the minor axis diameter. (C) Quantification of the mineralized bone volume (BV) in the *Pdpn* WT and KO E17.5 fetal femurs. BV was normalized to the total bone collar volume (TV). (D) Representative immunohistochemical images of the *Pdpn* WT and KO E17.5 fetal femurs. OSX and CD31 are shown in green and red, respectively. (E) Quantification of OSX-positive cells in the bone collars of *Pdpn* WT and KO E17.5 fetal femurs. Data are shown as cell density of OSX-positive cells in square micrometer. In panels B, C, and E, data are represented as the mean ± SD (n = 4 in *Pdpn* WT mice; n = 5 in *Pdpn* KO mice). Statistical comparisons were performed via Welch’s unpaired *t*-test. \**p* < 0.05, WT vs. KO. Scale bar, 100 μm.

**Figure 8.**
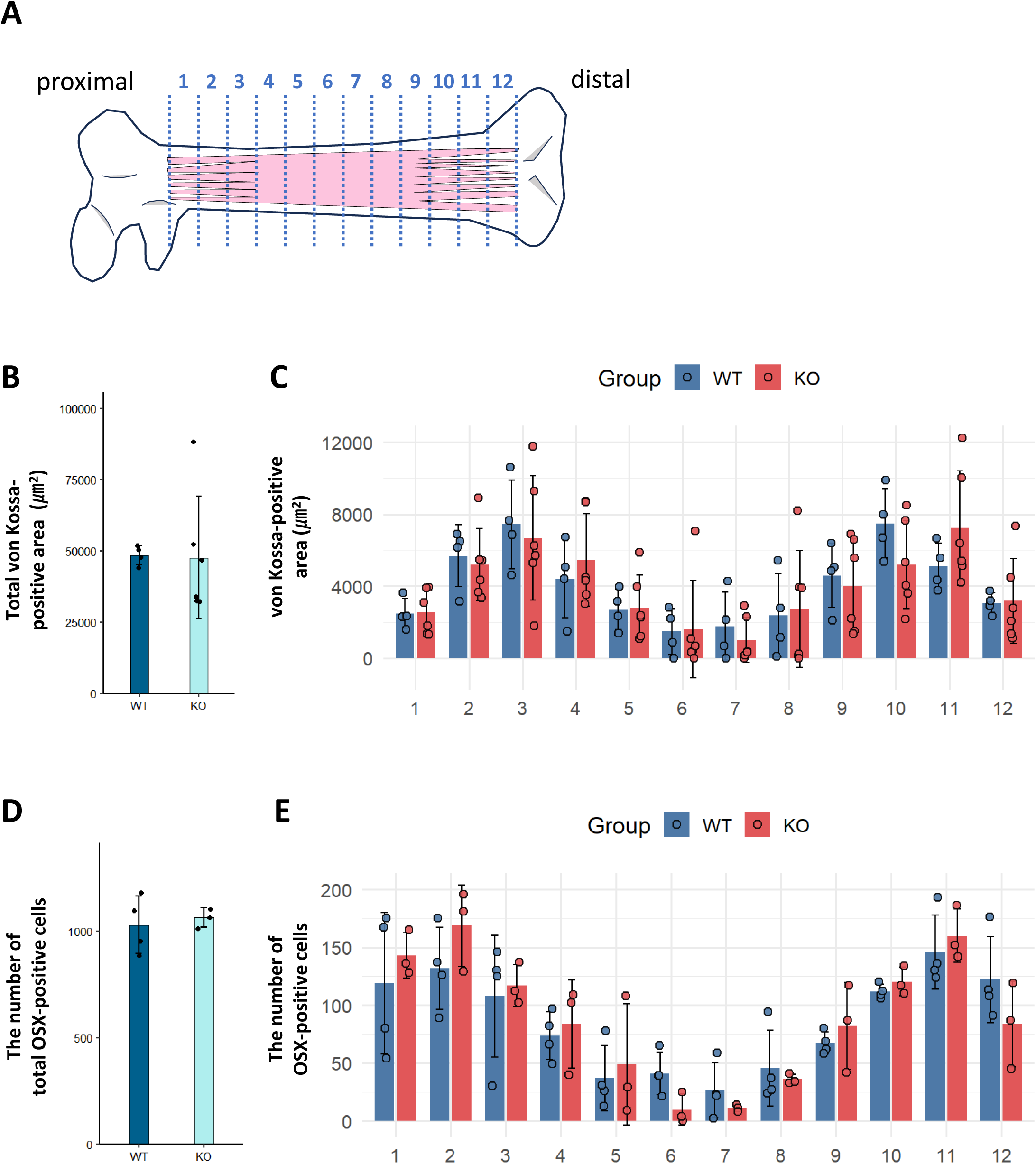
Osteogenesis of the trabecular bone in fetal *Pdpn* KO mice. (A) Schematic illustration of the 12 longitudinal divisions of the marrow cavity, including the trabecular regions. (B) Total von Kossa-positive area in the marrow cavity, including the trabecular regions. (C) Distribution of the von Kossa-positive area in each division. (D) Quantification of the total number of OSX-positive cells in the marrow cavity, including the trabecular regions. (E) Distribution of the number of total OSX-positive cells in each division. In panels B-E, data are represented as the mean ± SD (n = 4 in *Pdpn* WT mice; n = 5 in *Pdpn* KO mice). Statistical comparisons were performed via Welch’s unpaired *t*-test. \**p* < 0.05, WT vs. KO. Scale bar, 100 μm.

First, we focused on fetal bone collar formation (Fig. 7). Because the thickness of the femoral bone collar differs between the dorsal and ventral sides, we analyzed these regions separately. Bone collar was visualized via von Kossa staining, and its thickness was normalized to the minor axis diameter of the femur (Fig. 7A). Notably, no significant difference in femur diameter was observed between WT and KO mice (WT: 489.29 ± 16.09 μm; KO: 479.35 ± 7.53 μm; *p* = 0.6481, Welch’s unpaired *t*-test). Moreover, no significant difference in bone collar thickness was observed between the WT and KO mice on both the dorsal (WT: 0.317 ± 0.050; KO: 0.317 ± 0.027) and ventral (WT: 0.190 ± 0.045; KO: 0.201 ± 0.022) sides (Fig. 7B). However, mineralized bone volume (BV), normalized to the total bone collar volume (TV), was significantly reduced on the dorsal side in KO mice (WT: 30.7 ± 4.4%; KO: 22.6 ± 4.8%; *p* < 0.05, Welch’s unpaired *t*-test; Fig. 7C). BV/TV on the ventral side showed a decreasing trend but did not reach statistical significance (WT: 32.7 ± 5.9%; KO: 25.6 ± 7.0%; *p* = 0.13, Welch’s unpaired *t*-test). To assess osteogenic activity, we quantified the density of OSX-positive cells in the bone collar (Fig. 7D and E). On the dorsal side, OSX-positive cell density was significantly lower in KO mice than in WT mice (KO vs. WT: 0.287 ± 0.066 vs. 0.524 ± 0.066 cells/100 μm²; *p* < 0.01, Welch’s unpaired *t*-test; Fig. 7E, dorsal). Although a decreasing trend in OSX-positive cell density was also observed on the ventral side, the difference did not reach statistical significance (KO vs. WT: 0.398 ± 0.119 vs. 0.512 ± 0.076 cells/100 μm²; *p* = 0.24, Welch’s unpaired *t*-test; Fig. 7E, ventral). These findings suggest that the genetic deletion of *Pdpn* leads to impaired bone mineralization and reduced recruitment of OSX-positive osteogenic cells, particularly on the dorsal side of the bone collar.

We further examined trabecular bone formation (Fig. 8). To evaluate the trabecular architecture, we divided the marrow cavity, including the trabecular regions, longitudinally into 12 regions (Fig. 8A) and quantified the von Kossa-positive regions and number of OSX-positive cells in each region (Fig. 8B–E). Total von Kossa-positive area across all 12 regions did not significantly differ between the WT and KO mice (Fig. 8B). Distribution analysis revealed two peaks in regions 2–4 and 9–11, corresponding to the proximal and distal trabecular zones, respectively (Fig. 8C). However, no genotype-dependent differences were detected in the patterning and extent of trabecular mineralization. Similarly, total number and distribution of OSX-positive cells were comparable between both genotypes (Fig. 8D and E). Collectively, these results suggest that *Pdpn* deletion selectively impairs mineralization and osteogenic cell recruitment in the bone collar, particularly on the dorsal side, but is dispensable for initial trabecular bone formation.

## Discussion

In this study, we elucidated the spatiotemporal dynamics of PDPN-positive cells in the developing fetal femur and their physiological contribution to the bone collar microarchitecture. In the fetal femur, PDPN-positive cells first emerged in the bone collar concurrently with POC initiation. PDPN-positive cells in the bone collar expanded and subsequently migrated into the developing marrow cavity. During this transition, the cells resided in the marrow cavity-side of the trabecular bone. Most PDPN-positive cells residing in both the bone collar and trabecular regions exhibited osteolineage features, such as OSX expression. Genetic ablation of *Pdpn* led to the aberrant recruitment of OSX-positive cells and mineral deposition on the dorsal bone collar. These findings suggest that PDPN-positive cells constitute a spatially regulated osteolineage population contributing to coordinated fetal femur development.

Bone collar/cortical bone is a bone modeling site in which bone formation and resorption are spatially and temporally independent. Several studies have reported the association between PDPN expression and cortical bone modeling in adult mice ^15–20^. However, specific roles of PDPN in fetal bone development remain unclear. Systemic *Pdpn* KO mice exhibit perinatal lethality due to impaired lung development, specifically abnormally expanded alveolar sacs during late gestation ^24–28^. To date, only one study by Zhang et al. ^23^ has shown that femoral morphological parameters, such as femoral length, bone collar thickness, and diaphyseal minor diameter, in *Pdpn* KO fetuses are comparable to those in WT mice. Consistent with this report, *Pdpn* KO fetuses did not show significant differences in gross morphological indices in this study (Fig. 7). On E15.5, onset of bone collar formation following POC initiation was marked by the prominent invasion of OSX-positive and PDPN-negative osteoblasts into the cartilage matrix, with only some PDPN-positive cells present at this stage (Fig. 5). From E16.5 onward, two distinct OSX-positive osteoblast populations, PDPN-positive and PDPN-negative populations, were observed in the bone collar, suggesting that PDPN-positive cells constitute a subpopulation within the osteoblast lineage. These results suggest that OSX⁺/PDPN⁻ osteoblasts are the primary contributors to early bone collar formation, and OSX⁺/PDPN^+^ osteoblasts are involved in enhancing the quality of bone collar/cortical bone. Taken together, our findings suggest that the lack of significant morphological changes in the femurs of *Pdpn* KO mice reflects the limited involvement of PDPN-positive cells in the initial shaping of the bone collar.

On the other hand, our extended analysis revealed reduced mineral deposition in the bone collar, particularly on the dorsal side (Fig. 7). We consider that this low mineralization was due to a decrease in the number of OSX-positive osteoblasts in the bone collar. At the cellular level, PDPN possibly affects osteogenesis via three main mechanisms: (1) Differentiation of osteoblasts into osteocytes ^15,16,19,29^, (2) osteocytic dendrite formation ^17,19,23^, and (3) osteoblast migration ^16,30,31^. Moreover, Takenaka et al. suggested that PDPN plays a role in mineralization in cooperation with OCN and osteopontin ^32^. Genetic ablation of *Pdpn* possibly impairs the PDPN-mediated processes, including osteoblast migration, maturation, and mineralization, leading to reduced bone quality, particularly in the dorsal bone collar, without altering the overall bone morphology.

In the adult femur, metaphyseal trabecular bone is the primary site of bone remodeling, where bone formation and resorption occur in a spatially and temporally coordinated manner. Nagai et al. ^16^ reported that a subset of PDPN-positive osteoblasts in this region differentiate into osteocytes via interactions between PDPN and CD44. In this study, PDPN-positive cells resided in the marrow cavity-side region of the trabecular bone (Figs. 1 and 3). Therefore, we hypothesized that PDPN-positive cells support trabecular bone formation and maintenance. However, *Pdpn* KO fetuses exhibited no apparent abnormalities in the trabecular morphology and mineral content (Fig. 8). These findings suggest that PDPN is dispensable for trabecular bone formation during fetal development. Our extended investigation provided further insights into the cellular heterogeneity of PDPN-positive cells in the trabecular region. Specifically, we identified three immunophenotypic subpopulations: (1) PDPN⁺/OSX⁺/OCN⁺, (2) PDPN⁺/OSX⁺/OCN⁻, and (3) PDPN⁺/OSX⁻/OCN⁻ (Fig. 6). The first two groups belong to the osteogenic lineage, representing the different maturation stages of osteoblasts. However, identity and functional role of the third group, PDPN⁺/OSX⁻/OCN⁻, remain uncertain. In humans, PDPN acts as a molecular marker of skeletal stem/progenitor cells, a specialized subset of MSCs responsible for skeletal development, homeostasis, and regeneration ^33^. We previously demonstrated that PDPN-positive cells constitute a minor subset of skeletal stem/progenitor cells in mice ^34^. These PDPN⁺/OSX⁻/OCN⁻ cells possibly represent a pool of uncommitted progenitors poised for lineage commitment, such as the osteogenic lineage. Although the mechanisms by which the PDPN-positive cells restrictedly localize to the marrow cavity-side of the trabeculae were unclear, we speculated that the PDPN-positive cells, including the population with PDPN⁺/OSX⁻/OCN⁻ cells, contribute to postnatal bone remodeling in the metaphyseal region, particularly during longitudinal bone growth. Further cellular identification of the PDPN⁺/OSX⁻/OCN⁻ cells will provide key insights into metaphyseal bone remodeling.

The above-mentioned findings raise questions regarding the sites at which PDPN-positive cells bind to their ligands during bone and marrow development. PDPN has eight types of protein partners ^35^. C-type lectin-like receptor 2 (CLEC-2) is the major PDPN ligand ^28^. In long bones, CLEC-2 is expressed in the hematopoietic cells, such as the long-term hematopoietic stem cells ^36,37^ and megakaryocytes/platelets ^38–41^. We examined whether PDPN-positive stromal cells spatially co-localize with CLEC-2– positive hematopoietic cells in the developing bone and marrow; however, no direct contact between these cells was observed (data not shown). Another important ligand, CD44, forms the PDPN/CD44 axis, which promotes the migration and differentiation of PDPN-positive osteogenic cells ^16,30,31^. CD44 is expressed in the bone marrow cells, osteoclasts, and osteoblasts of adult femurs ^16^. In the developing fetal long bone, we consider that PDPN-positive cells in the bone collar can encounter CD44-expressing cells, thereby receiving pro-migratory and pro-differentiation cues via the PDPN/CD44 axis. To clarify the molecular and cellular functions of PDPN-positive cells in fetal bone formation, particularly bone collar formation, future studies using PDPN/CD44 axis-targeted genetically engineered model mice are necessary to elucidate the mechanisms by which PDPN/CD44 signaling spatially regulates osteogenic differentiation and stromal cell positioning during fetal ossification.

In conclusion, this study revealed that PDPN-positive cells emerging from the osteogenic lineage played roles in fetal bone development. However, this study has some limitations. Perinatal lethality of *Pdpn*-KO mice restricted the investigation to only embryonic and fetal stages. The mechanisms by which the reduced quality of the bone collar during the fetal stage affects postnatal cortical bone development remain unclear. Further investigations using highly efficient and lineage-restricted mouse models, such as the *Sp7*-cre;*Pdpn* conditional KO model or acquired and site-specific PDPN-positive cell depletion model, are essential to clarify the PDPN-positive cell functions during postnatal bone development. Additional studies are pivotal to elucidating the ontogenic and functional roles of PDPN-positive cells in bone formation, maintenance, and repair.

## Acknowledgments

We are grateful to the Nikon Imaging Center at Hokkaido University for assistance with confocal microscopy, image acquisition, and analysis. We would like to thank Editage (www.editage.jp) for English language editing.

## CRediT author contribution statement

H.N., N.T., and S.T., Methodology; H.N., N.T., and S.T., Investigation; H.N. and S.T., Formal analysis; H.N. and S.T., Writing – original draft; N.T., N.S., A.S., S.O., T. Kanematsu, A. Katsumi., R.N., Ayuka K., T. Kojima, T.M., and K.S-I., Supervision; S.T., Conceptualization; S. T., Writing – review & editing; H.N. and S.T., Data curation; Nobuaki S., A. Katsumi., T.M., and S.T., Funding acquisition; S.T., Validation; S.T., Visualization; S.T., Project administration.

## Materials and Methods

### Mice

C57BL/6NJc1 mice were purchased from CLEA Japan (Tokyo, Japan). *Pdpn* KO mice were generated as previously described ^27^. All mice were maintained under a 12/12-h light/dark cycle under standard conditions, with controlled temperature (25 °C), and humidity (60%). All animal experiments were approved by the Animal Care and Use Committee of the Hokkaido University (22-0087) and University of Yamanashi, Japan (A29-49 and A3-61).

### Tissue section preparation

Femur was fixed with 4% paraformaldehyde (PFA; Wako, Japan) for 24 h and subsequently decalcified in K-CX solution (FALMA, Tokyo, Japan) for 3–5 d depending on the mouse age. After thoroughly washing with phosphate-buffered saline (PBS), the samples were incubated with 30% sucrose in PBS for 24 h for cryoprotection. The solution was sequentially replaced with a 1:1 mixture of 15% sucrose in PBS and the Tissue-Tek O.C.T. Compound (Sakura Finetek, Tokyo, Japan), followed by the 100% Tissue-Tek O.C.T. Compound. The tissues were embedded in the O.C.T. compound and frozen at −80 °C. Subsequently, cryosections were cut to a thickness of 10 μm using a cryostat (Leica CM1950, Tokyo, Japan).

### IHC staining

Frozen sections were blocked with Blocking One Histo (Nacalai Tesque, Kyoto, Japan) for 7 min and washed with 0.05% PBS-Tween 20 (PBS-T) for 10 min. Fc receptor was blocked using the rat anti-mouse CD16/CD32 antibodies (1:200 dilution in antibody diluent; BD Biosciences, NJ, USA) for 30 min. The antibody diluent consisted of Blocking One Histo (diluted 1:20 in PBS-T.) After washing twice with PBS-T for 10 min each, the sections were incubated with primary antibodies overnight at 4 °C. The following day, the sections were washed thrice with PBS-T and incubated with the appropriate secondary antibodies for 1.5 h at room temperature. All antibodies used in this study are listed in Supplementary Table S1. For double immunostaining using goat polyclonal anti-CD31 and hamster monoclonal anti-PDPN antibodies, both primary antibodies were simultaneously applied. Then, the sections were sequentially incubated with the fluorescein-conjugated donkey anti-goat IgG and goat anti-hamster IgG secondary antibodies. The sections were mounted with the VECTASHIELD Antifade Mounting Medium containing 4’,6-diamidino-2-phenylindole (DAPI) (Vector Laboratories, CA, USA) and imaged using an upright fluorescence microscope (ECRIPSE E600, Nikon, Tokyo, Japan) equipped with the Plan Fluor ×10 objective lens (NA 0.30; Nikon), Plan Fluor ×20 objective lens (NA 0.50; Nikon), and Plan Fluor ×40 objective lens (NA 0.75; Nikon) and inverted confocal laser scanning microscope (A1Rsi; Nikon) equipped with the Plan Fluor x40 objective lens (NA 0.75; Nikon). The acquired images were analyzed using the ImageJ software (version 1.46r; http://rsb.info.nih.gov/ij/).

### HE, alcian blue, and von Kossa staining

Frozen sections were post-fixed with fixative solutions composed of ethanol, formaldehyde, and acetic acid for 5 min, followed by washing with purified water. For HE staining, the sections were stained with Mayer’s hematoxylin solution (Wako) for 5 min and rinsed thoroughly with running water to develop color. Subsequently, eosin solution (Wako) was applied for 1 min. For alcian blue staining, frozen sections were stained with the alcian blue solution (Wako) for 10 min. As a counterstain, fast red solution (ScyTek Laboratories, UT, USA) was applied for 1 min. After rinsing with purified water, the sections were dehydrated using a graded ethanol series (50, 70, 100, and 100%), cleared with Lemosol (Wako), and mounted using a permanent mounting medium (Multi Mount 480; Matsunami Glass, Tokyo, Japan). For von Kossa staining, frozen sections were post-fixed with 4% paraformaldehyde (Wako) for 5 min and washed with distilled water twice for 10 min. Calcium deposition was visualized using the Calcium Stain Kit (ScyTek Laboratories), according to the manufacturer’s instructions. Nuclei were counterstained with the fast red solution.

### Quantitative image data analysis (area measurement and cell count)

Number of PDPN-positive cells in the cortical bone was quantified in the transverse (short-axis) sections of the femur, and that in the metaphysis was quantified in the longitudinal (long-axis) sections of the femur (Fig. 4). For each sample, three representative sections were analyzed, and the average value was calculated. Cell counts were normalized to the areas of the respective regions. Numbers of OSX-positive cells in the bone collar (Fig. 7) and trabecular bone regions (Fig. 8) were evaluated in the longitudinal (long-axis) sections on E17.5. For each sample, two sections were analyzed, and the mean was calculated. Cell counts were normalized to the corresponding tissue areas. Bone collar thickness and bone volume fraction (BV/TV; bone volume/total volume, %) were calculated from the von Kossa-stained sections (Fig. 7). Bone collar thickness was normalized to the width of the femoral diaphysis.

For each sample, two sections were analyzed, and the mean value was calculated. To quantify the mineralized area in the trabecular bone, von Kossa-positive area was measured and expressed as a percentage of the total tissue area (Fig. 8). To evaluate the structural abnormalities in the trabecular bone, bone marrow cavity was divided into 12 equally spaced regions, from proximal to distal, and the von Kossa-positive area was quantified in each region (Fig. 8). Image analysis was performed using the ImageJ software.

### Statistical analyses

Quantitative data are represented as the mean ± standard deviation. One-way analysis of variance (ANOVA) followed by the Tukey’s post-hoc test, was used for multigroup comparisons. Statistical comparisons between two groups were performed via Welch’s unpaired *t-*test. All statistical analyses were conducted using R (version 4.5.0) and RStudio.

## References

1 Kronenberg, H. M. Developmental regulation of the growth plate. Nature 423, 332–336, doi:10.1038/nature01657 (2003).

2 Yang, L., Tsang, K. Y., Tang, H. C., Chan, D. & Cheah, K. S. Hypertrophic chondrocytes can become osteoblasts and osteocytes in endochondral bone formation. Proc Natl Acad Sci U S A 111, 12097–12102, doi:10.1073/pnas.1302703111 (2014).

3 Karsenty, G. & Wagner, E. F. Reaching a genetic and molecular understanding of skeletal development. Dev Cell 2, 389–406, doi:10.1016/s1534-5807(02)00157-0 (2002).

4 Karsenty, G., Kronenberg, H. M. & Settembre, C. Genetic control of bone formation. Annu Rev Cell Dev Biol 25, 629–648, doi:10.1146/annurev.cellbio.042308.113308 (2009).

5 Maes, C. et al. Osteoblast precursors, but not mature osteoblasts, move into developing and fractured bones along with invading blood vessels. Dev Cell 19, 329–344, doi:10.1016/j.devcel.2010.07.010 (2010).

6 Crivellato, E. The role of angiogenic growth factors in organogenesis. Int J Dev Biol 55, 365–375, doi:10.1387/ijdb.103214ec (2011).

7 Matsushita, Y. et al. The fate of early perichondrial cells in developing bones. Nat Commun 13, 7319, doi:10.1038/s41467-022-34804-6 (2022).

8 Trompet, D., Melis, S., Chagin, A. S. & Maes, C. Skeletal stem and progenitor cells in bone development and repair. J Bone Miner Res 39, 633–654, doi:10.1093/jbmr/zjae069 (2024).

9 Maes, C., Kobayashi, T. & Kronenberg, H. M. A novel transgenic mouse model to study the osteoblast lineage in vivo. Ann N Y Acad Sci 1116, 149–164, doi:10.1196/annals.1402.060 (2007).

10 Breiteneder-Geleff, S. et al. Podoplanin, novel 43-kd membrane protein of glomerular epithelial cells, is down-regulated in puromycin nephrosis. Am J Pathol 151, 1141–1152 (1997).

11 Wetterwald, A. et al. Characterization and cloning of the E11 antigen, a marker expressed by rat osteoblasts and osteocytes. Bone 18, 125–132, doi:10.1016/8756-3282(95)00457-2 (1996).

12 Farr, A. G. et al. Characterization and cloning of a novel glycoprotein expressed by stromal cells in T-dependent areas of peripheral lymphoid tissues. J Exp Med 176, 1477–1482, doi:10.1084/jem.176.5.1477 (1992).

13 Rishi, A. K. et al. Cloning, characterization, and development expression of a rat lung alveolar type I cell gene in embryonic endodermal and neural derivatives. Dev Biol 167, 294–306, doi:10.1006/dbio.1995.1024 (1995).

14 Cheok, Y. Y. et al. Podoplanin and its multifaceted roles in mammalian developmental program. Cells Dev 180, 203943, doi:10.1016/j.cdev.2024.203943 (2024).

15 Staines, K. A. et al. E11/Podoplanin Protein Stabilization Through Inhibition of the Proteasome Promotes Osteocyte Differentiation in Murine in Vitro Models. J Cell Physiol 231, 1392–1404, doi:10.1002/jcp.25282 (2016).

16 Nagai, T. et al. Immunocytochemical assessment of cell differentiation of podoplanin-positive osteoblasts into osteocytes in murine bone. Histochem Cell Biol 155, 369–380, doi:10.1007/s00418-020-01937-y (2021).

17 Staines, K. A. et al. Hypomorphic conditional deletion of E11/Podoplanin reveals a role in osteocyte dendrite elongation. J Cell Physiol 232, 3006–3019, doi:10.1002/jcp.25999 (2017).

18 Takara, K. et al. Morphological study of tooth development in podoplanin-deficient mice. PLoS One 12, e0171912, doi:10.1371/journal.pone.0171912 (2017).

19 Ikpegbu, E. et al. FGF-2 promotes osteocyte differentiation through increased E11/podoplanin expression. J Cell Physiol 233, 5334–5347, doi:10.1002/jcp.26345 (2018).

20 Kanai, T., Osawa, K., Kajiwara, K., Sato, Y. & Sawa, Y. Study of Podoplanin-Deficient Mouse Bone with Mechanical Stress. Dent J (Basel) 13, doi:10.3390/dj13020061 (2025).

21 Nefussi, J. R., Sautier, J. M., Nicolas, V. & Forest, N. How osteoblasts become osteocytes: a decreasing matrix forming process. J Biol Buccale 19, 75–82 (1991).

22 Barragan-Adjemian, C. et al. Mechanism by which MLO-A5 late osteoblasts/early osteocytes mineralize in culture: similarities with mineralization of lamellar bone. Calcif Tissue Int 79, 340–353, doi:10.1007/s00223-006-0107-2 (2006).

23 Zhang, K. et al. E11/gp38 selective expression in osteocytes: regulation by mechanical strain and role in dendrite elongation. Mol Cell Biol 26, 4539–4552, doi:10.1128/MCB.02120-05 (2006).

24 Ramirez, M. I. et al. T1alpha, a lung type I cell differentiation gene, is required for normal lung cell proliferation and alveolus formation at birth. Dev Biol 256, 61–72, doi:10.1016/s0012-1606(02)00098-2 (2003).

25 Schacht, V. et al. T1alpha/podoplanin deficiency disrupts normal lymphatic vasculature formation and causes lymphedema. EMBO J 22, 3546–3556, doi:10.1093/emboj/cdg342 (2003).

26 Millien, G. et al. Alterations in gene expression in T1 alpha null lung: a model of deficient alveolar sac development. BMC Dev Biol 6, 35, doi:10.1186/1471-213X-6-35 (2006).

27 Tsukiji, N. et al. Platelets play an essential role in murine lung development through Clec-2/podoplanin interaction. Blood 132, 1167–1179, doi:10.1182/blood-2017-12-823369 (2018).

28 Suzuki-Inoue, K. & Tsukiji, N. A role of platelet C-type lectin-like receptor-2 and its ligand podoplanin in vascular biology. Curr Opin Hematol 31, 130–139, doi:10.1097/MOH.0000000000000805 (2024).

29 Prideaux, M., Loveridge, N., Pitsillides, A. A. & Farquharson, C. Extracellular matrix mineralization promotes E11/gp38 glycoprotein expression and drives osteocytic differentiation. PLoS One 7, e36786, doi:10.1371/journal.pone.0036786 (2012).

30 Martin-Villar, E. et al. Podoplanin binds ERM proteins to activate RhoA and promote epithelial-mesenchymal transition. J Cell Sci 119, 4541–4553, doi:10.1242/jcs.03218 (2006).

31 Martin-Villar, E. et al. Podoplanin associates with CD44 to promote directional cell migration. Mol Biol Cell 21, 4387–4399, doi:10.1091/mbc.E10-06-0489 (2010).

32 Takenawa, T. et al. Expression and Dynamics of Podoplanin in Cultured Osteoblasts with Mechanostress and Mineralization Stimulus. Acta Histochem Cytochem 51, 41–52, doi:10.1267/ahc.17031 (2018).

33 Chan, C. K. F. et al. Identification of the Human Skeletal Stem Cell. Cell 175, 43–56 e21, doi:10.1016/j.cell.2018.07.029 (2018).

34 Tamura, S. et al. Periosteum-derived podoplanin-expressing stromal cells regulate nascent vascularization during epiphyseal marrow development. J Biol Chem 298, 101833, doi:10.1016/j.jbc.2022.101833 (2022).

35 Quintanilla, M., Montero-Montero, L., Renart, J. & Martin-Villar, E. Podoplanin in Inflammation and Cancer. Int J Mol Sci 20, doi:10.3390/ijms20030707 (2019).

36 Nakamura-Ishizu, A., Takubo, K., Kobayashi, H., Suzuki-Inoue, K. & Suda, T. CLEC-2 in megakaryocytes is critical for maintenance of hematopoietic stem cells in the bone marrow. J Exp Med 212, 2133–2146, doi:10.1084/jem.20150057 (2015).

37 Kumode, T. et al. C-type lectin-like receptor 2 specifies a functionally distinct subpopulation within phenotypically defined hematopoietic stem cell population that contribute to emergent megakaryopoiesis. Int J Hematol 115, 310–321, doi:10.1007/s12185-021-03220-9 (2022).

38 Suzuki-Inoue, K. et al. A novel Syk-dependent mechanism of platelet activation by the C-type lectin receptor CLEC-2. Blood 107, 542–549, doi:10.1182/blood-2005-05-1994 (2006).

39 Suzuki-Inoue, K. et al. Involvement of the snake toxin receptor CLEC-2, in podoplanin-mediated platelet activation, by cancer cells. J Biol Chem 282, 25993–26001, doi:10.1074/jbc.M702327200 (2007).

40 Suzuki-Inoue, K. et al. Essential in vivo roles of the C-type lectin receptor CLEC-2: embryonic/neonatal lethality of CLEC-2-deficient mice by blood/lymphatic misconnections and impaired thrombus formation of CLEC-2-deficient platelets. J Biol Chem 285, 24494–24507, doi:10.1074/jbc.M110.130575 (2010).

41 Osada, M. et al. Platelet activation receptor CLEC-2 regulates blood/lymphatic vessel separation by inhibiting proliferation, migration, and tube formation of lymphatic endothelial cells. J Biol Chem 287, 22241–22252, doi:10.1074/jbc.M111.329987 (2012).

